# Pollinator Species Richness Trends: No Decelerating Declines in Dutch Bees and Bumblebees

**DOI:** 10.1101/240432

**Authors:** Tom J. M. Van Dooren

**Author notes:** Statement of authorship: TJMVD designed the study, carried out the analysis and wrote the manuscript. Data accessibility statement: The raw data used are available upon request from http://www.eis-nederland.nl.

## Abstract

Temporal trends (1946–2013) in the species richness of wild bees from the Netherlands are analysed. We apply two methods to estimate richness change which both incorporate models for sampling effects and detection probability. The analysis is repeated for records with specimens deposited in collections, and a subset restricted to spatial grid cells that have been sampled repeatedly across three periods. When fitting non-linear species accumulation curves to species numbers, declines are inferred for bumblebees and at most limited declines for other bees. Capture-recapture analysis applied to species encounter histories infers a constant colonization rate per year and constant (bumblebees) or decreasing (other) local species survival. However, simulations suggest that the method estimates time trends in survival with a negative bias. Species richness trends predicted by the second approach are a 10% reduction in non-*Bombus* species richness and 29% fewer *Bombus* species since 1946, comparable to the predictions of the first approach. Neither analysis provides reliable evidence that decelerating declines in species richness occur in these taxa. Therefore we should not infer decelerating declines in pollinator species richness in N-W Europe as previously claimed.

## Introduction

Species richness, the number of species present in a community or assemblage, is an important component of biodiversity. Species richness trends of pollinators providing essential ecosystem services merit a lot of attention and different drivers of pollinator abundance and richness change have been identified (Potts et al. 2016a). While it has been shown that visitation to crop flowers increases with the number of species per field (Garibaldi et al. 2013), species richness might be not the key quantity predicting crop pollination services to agriculture, as common species provide most of these (Winfree et al. 2015). Kleijn et al. (2015) therefore state that conservation and immediate utility goals for agriculture might not align. However, in the face of climate change and when accounting for geographical crop variation, the conservation of pollinator richness levels might be crucial to mitigate effects of ecosystem change (Klein et al. 2009; Gonzales-Varo et al. 2013). It also seems that an overall and substantial decline in total biomass, such as observed for insects in nature reserves in Germany (Hallmann et al. 2017), will reduce species richness inevitably. In a recent analysis of species richness trends, Carvalheiro et al. (2013) concluded that declines in species richness have slowed down for several taxa in NW-Europe. A reassessment of that analysis (Van Dooren 2016) concluded that it only provided support for decelerating declines for the bees *Anthophila* in the Netherlands, conditional on accepting the inference method and parameter estimates as valid.

Carvalheiro et al. (2013) rightly stated that detecting decelerations is highly relevant for conservation biology and biodiversity management. However, that should occur while ensuring conservative statistical inference, to avoid spurious conclusions with management consequences that might be difficult to repair. Conservative here refers to a concept of statistical inference, which implies that we keep the amount of type I errors below nominal level, even when sacrificing statistical power. It also has a general ethical meaning that applies here: we should be conservative in our inference not to accidentally increase extinction risks.

The estimation of species richness has been given a lot of attention already (Gotelli & Colwell 2011), that of species richness change over time less so. Species richness is the horizontal asymptote of a species accumulation curve (the expected number of species sampled as a function of sample size) and estimators are often biased (Walther & Moore 2005), or the precision of an estimate can be limited (O’Hara 2005). In a comparison of two assemblages, species accumulation curves can cross (Lande et al. 2000; Thompson & Withers 2003; Chao & Jost 2012; Van Dooren 2016), such that patterns in the relative numbers of species found at low sampling efforts can be independent of actual species richness differences. For example, the highly cited report by Biesmeijer et al. (2006) on pollinator decline interpolates the data to the lowest sampling efforts of pairs of samples (“rarefaction”). Without supporting evidence that species accumulation curves do not cross, such an analysis allows only weak inference on true relative species richness. Inference on parameters representing changes in species richness might proceed differently from the estimation of species richness itself and patterns of bias and precision do not need to be the same in these different parameterizations. A relevant case is found in the difference in estimation bias between estimates of population size and of individual survival between two time points (Cormack 1972). Survival is a parameter determining changes in population size and is generally much less biased than population size estimators.

Next to the general importance of investigating time trends in species richness, there are several methodological arguments for reanalysing the Dutch bee data in Carvalheiro et al. (2013). There is general scepticism towards estimation of richness using non-parametric estimates (O’Hara 2005). Inference based on them does still improve (Chao & Jost 2012) with seemingly decreased bias for comparisons made at a fixed sample completeness (fraction of individuals in the assemblage belonging to species represented in the sample). However, there is no general strategy for comparing more than two samples or for estimating a time trend. In addition, the Dutch wild bee data are a heterogeneous mix: the records (sensu Isaac & Pocock 2015) were collected in different ways (observations, samples deposited in collections, by hundreds of observers and collectors) and relatively unplanned. This might be exemplary for most data collected in citizen science efforts but was unaccounted for. In the analyses of the data thus far, data were binned in arbitrary time periods while a year-to-year continuous time analysis would be insightful and could provide more detail on patterns of change. Van Dooren (2016) found a correct assessment of the analysis in Carvalheiro et al. (2013) difficult due to the absence of statistics and the presence of uncorrected errors. Here, for the re-analysis, I chose to reconsider the inference strategy. It applies two approaches for assessing species richness trends among four explored (Supplementary Material) and compares their merits, assisted by simulations.

## Material and Methods

Bee records in the EIS (European Invertebrate Survey Netherlands) database in the period 1946–2013 are analysed, extending periods investigated in Biesmeijer et al. (2006) and Carvalheiro et al. (2013). Sampling has not occurred in a standardized manner across the study period (Figure 1). In particular from mid-eighties until around 2005, increasingly larger numbers of records were collected, more often by observation and with records from a much larger number of 10×10 km grid cells. The samples show important time trends in the heterogeneity parameter a of Poisson lognormal distributions when fitted to the numbers of records per species in each year (Engen et al. 2002). Here, this parameter σ is a compound measure representing abundance variation between species rescaled by species-specific sampling effort if present. Temporal patterns in estimated species richnesses are expected to reflect these changes in sampling strategy on top of the actual biodiversity changes. I assessed patterns of species richness change over time in the entire Netherlands. To facilitate comparisons with Carvalheiro et al (2013), time patterns in species richness of *Bombus* and non-*Bombus* bee genera in the EIS database were analysed separately and in different ways, which each tried to take changes in sampling effort and method over time into account. Each of these approaches should be seen as an attempt to arrive at a satisfactory model for richness change, where the inference is kept conservative with respect to claims of richness recovery, i.e., a rejection of the null hypothesis of no deceleration in richness decline should not be spurious. Within each approach, model simplification is carried out with the purpose of obtaining the best possible predictions, henec by means of AICc comparisons. Model selection, parameter estimation and inference in general were backed up as follows. Next to an analysis of all data, two subsets were analysed separately and for each method: 1) Records for which the individual was deposited in a museum collection, excluding for example records based on observations only. 2) Records from 10×10 km grid cells that were sampled in the period where the number of observations drastically increased (1993–2013), in a period of equal length right after WWII (1946–1965) and in the years in between (1966–1992). In this manner, spatial locations that contributed samples in a restricted time period and with potentially high leverage on time trend estimates were excluded. Model simplification increases the precision of individual parameter estimates, but this can come at the cost of larger estimation bias. In each statistical analysis presented below, I assessed effects of this bias-variance trade-off (Claeskens & Hjort 2008) by comparing the predicted time trend of species richness between the best maximal model fitted and a simplified adequate model (see below). I will only conclude that a deceleration occurs when it is detected in the full data and the data subsets, and does not suddenly emerge as a result of model selection bias. To understand patterns of estimation bias and power, the analysis was further complemented with simulations (Supplementary Material).

**Figure 1.**
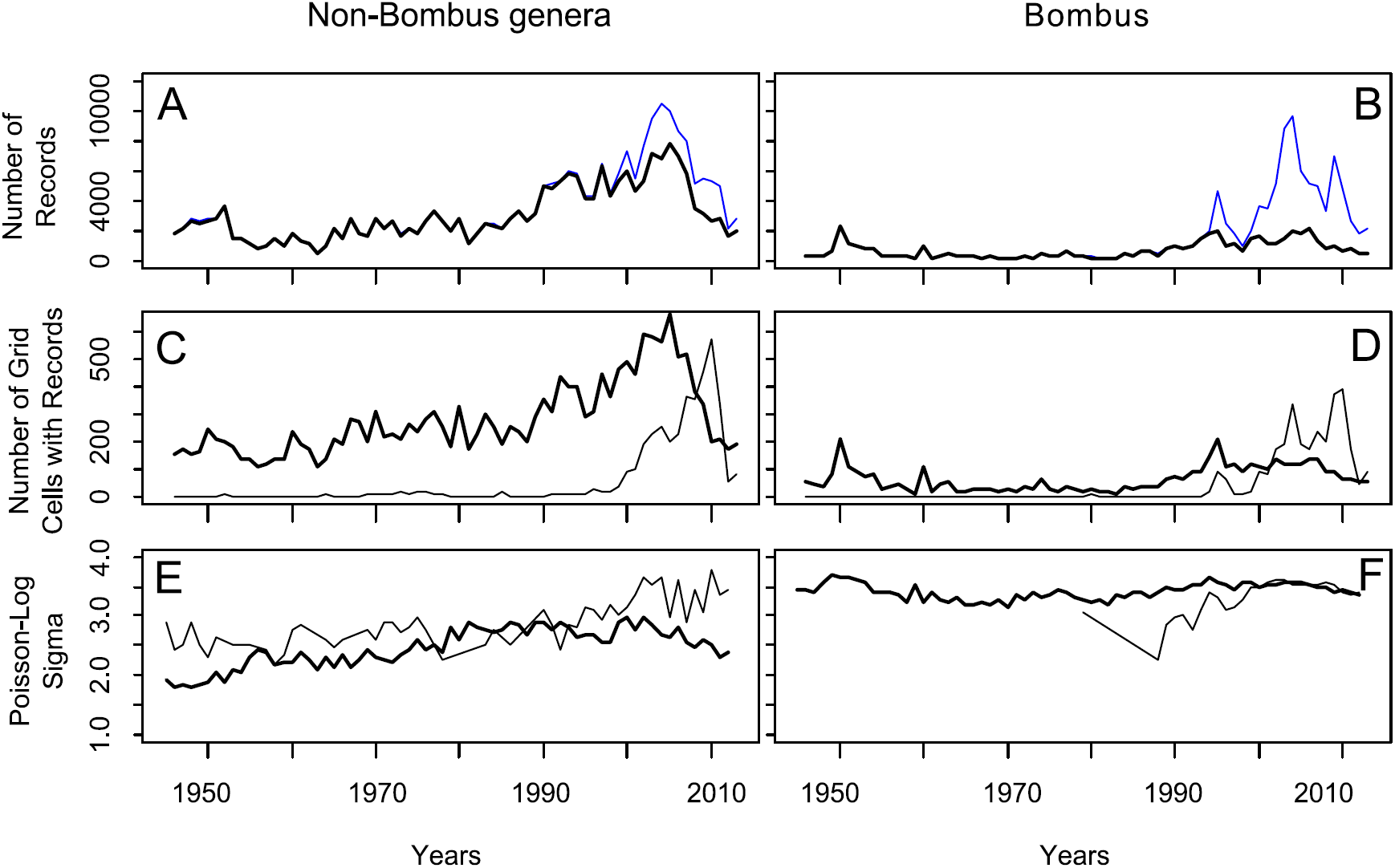
Sampling strategy and volume of wild bee data in the Netherlands have changed over the years. Top row. Total numbers of records have changed substantially (thin blue lines) and the fraction of records that represent individuals deposited in collections as well (thick lines, number of records with samples in collections). Middle row. Records are obtained in a variable number of 10km grid cells per year (thick: samples deposited in collections, thin: observations only). Bottom row. When distributions of records per species per year are analysed using Poisson log-normal models (Engen et al. 2002), the *σ* heterogeneity parameter for their distribution shows a gradual change over the years (thick: samples deposited in collections, thin: observations).

## Generalized non-linear models

In a first approach, the number of records and the number of species per year were used to estimate non-linear species accumulation curves (Gotelli & Colwelll 2011, Supplementary Material) and their changes over time. Species accumulation curves were represented by Michaelis-Menten curves *a*(*t*)*x*(*t*)*/*(*b*(*t*)*+x*(*t*)) estimating species number with two year-dependent non-negative functions *a*(*t*) and *b*(*t*) and with *x*(*t*) the number of records in year *t*, representing sample size. I assumed that functions *a*(*t*) and *b*(*t*) change gradually across years and therefore used smooth functions (Wood 2006) to parameterize them and assess their gradual changes. We can expect that *b*, which is determined by the increase in species number with sampling effort shows a time pattern affected by changes in sampling method, while *b/a* is an estimate of the Simpson diversity in random samples (Lande et al. 2000). If species richness, given by the asymptote *a*, has a decelerating decline, we should be able to observe that in the predicted time pattern for *a*(*t*). Note that this model representing a species accumulation curve combines the estimation of state variable *a*(*t*) with properties of the observation process captured by *b*(*t*).

For the fitting of Michaelis-Menten curves, I used the function *gnlr*() in R (Lindsey 1997) which maximizes the model likelihood using a general purpose optimizer. Functions *a* and *b* in the model were fitted with natural cubic splines of the year variable of up to eight degrees of freedom and exponential link functions assuring non-negativity of *a* and *b*. Normal errors were assumed (Colwell et al. 2012) and the error variance consisted of the sum of an estimated variance parameter plus an offset equal to the unconditional sampling variance of the number of observed species given multinomial sampling of individuals (eq. 5 in Colwell et al. 2012). This last variance term differed between years and the variance regression equation ensured that the error variance never became smaller than this multinomial sampling variance (Supplementary Material). For the curvature function *b*(*t*), alternative models with the same number of parameters but splines of the total number of grid cells with data per year or of the *σ* parameter of the Poisson lognormal distribution (Engen et al. 2002) fitted to the data per year for the group analysed were also tried. Model simplification occurred in a manner which slightly reduced the total number of models to fit and compare. I fitted all models with splines of 8, 5, 3 and 1 degrees of freedom and the models with no explanatory variables for *a* or *b*. Among the models in this set, I selected the one with the lowest AICc (Akaike information Criterion adjusted for finite sample sizes, Burnham & Anderson 2002). Then models with one d.f. added to each spline in this model or with one d.f. removed were also fitted, to check whether these modifications would reduce the AICc further. The model with the lowest AICc among all the ones fitted is called the minimum “adequate” model. This model is compared with the “maximal” model: the model with lowest AICc among those with splines of 8 d.f. for both *a*(*t*) and *b*(*t*).

To assess estimation bias, a corrected analysis was carried out, using a bootstrap estimate of bias (Davison & Hinkley 1997). One hundred bootstrap pseudo-datasets were constructed by randomly drawing for each year a number of individuals equal to the number of records in that year and repeating the analysis above for each of these sets of pseudo-data. The covariates calculated from the original data were kept except for the multinomial sampling variances which were re-calculated for each sample. The maximal and adequate models were fitted to each of these datasets, and species richnesses per year predicted. The estimated bootstrap bias equalled the average of these predictions per year minus the predicted value obtained from the original data. Note that this straightforward bootstrap differs from the one proposed by Chao et al. 2014, in that the number of undetected species is not estimated and incorporated in the resampling and that it is not used to obtain an unconditional estimate of sampling variance.

### Capture-recapture analysis of species encounter histories

Suggestions that methods of the capture-recapture framework should be applicable to assemblages of species encounter histories have been made repeatedly (Nichols & Pollock 1983; Nichols et al. 1998; MacKenzie et al. 2006). In this framework, every species encounter history is a sequence of zeroes and ones indicating for which years at least one record is present in the dataset. However, the dataset and the subsets analysed here have no repeated within-year occasions where the assemblage can be assumed to be closed, such that a robust design model cannot be fitted (Pollock 1982) and temporary emigration cannot be estimated in that manner. Temporary emigration can also be estimated assuming an unobservable state (Kendall & Nichols 2002, Schaub et al. 2004). Such a model as constructed for capture histories of individuals cannot deal with individual species jointly present inside and outside of the Netherlands and we would have to use occupancy models. For parameter estimability, we would further have to assume the absence of time trends in emigration and survival (Schaub et al. 2004) such that the time-dependent models of interest cannot be fitted. Therefore, encounter histories per species were analysed using capture-mark-recapture analysis for open populations (Pradel 1996) and trends in temporary emigration were assessed otherwise. Models were fitted to species encounter histories across years using Mark software (White & Burnham 1999) called via RMark in R (Laake 2013). I parameterized using the “Pradrec” model, which estimates local survival and colonization, where I note that the first is a probability per species (in between zero and one) and the second the number of colonizing species relative to the number present at the beginning of a time interval (non-negative). Local survival probabilities, species colonization and detection probabilities were estimated as being constant, with a linear trend over time (in the linear predictor), with categorical effects per year (“time-dependent”) or with regression models that have the total number of records, number of grid cells with records per year, and *σ* of the Poisson lognormal distribution of the taxon concerned as year-specific covariates (“regression”). Correlated explanatory variables were fitted in the detection probability model because the aim was to obtain the best predictions for survival and recruitment, not to interpret parameter estimates of the detection model or do hypothesis testing on them. The Pradel models fit parameters that express per-year gains and losses in species richness and not richness itself. This is expected to reduce estimation bias (Cormack 1972) but it is also known that capture heterogeneity needs to be addressed (Abadi et al. 2013) to avoid biased estimates of local survival and colonization. For that reason, mixtures of species effects on detection probability were fitted, with two or three components (Pradel et al. 2009). Models were compared using AICc to select the minimum adequate model with lowest AICc. AIC(c) model selection is efficient (Claeskens & Hjort 2008). If the probability that a species is temporarily absent from the population is constant over time, it will not affect estimates of species richness change. The presence of time-inhomogeneous temporary emigration could affect time trends in species richness and was assessed as follows. If the probability of being temporarily absent in year *t* is *p_e_*(*t*) and the probability of detection when present *p_d_*(*t*), then the probability of detection *p*(*t*) in the time-dependent Pradel model should equal the product of *p_d_*(*t*) and (1-*p_e_*(*t*)). Assuming that our regression model for detection probabilities with variables characterizing sampling effort and method fits in fact *p_d_*(*t*) multiplied by a constant, we can investigate the presence of elevated temporary emigration in some time intervals by checking time patterns of the difference between detection probabilities predicted by the regression and by the fully time-dependent models. Large discrepancies in a sequence of consecutive years can indicate temporary emigration affecting species richness trends.

Parameter estimates of local survival and colonization were used to calculate the change in species number in 2017 relative to 1945. The growth rate *λ_t_* of species richness in between year *t* and *t* + 1 is the sum of the survival probability *s*_t_ and colonization *f*_t_. The predicted relative change in richness in year 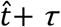 relative to reference year 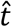 is equal to

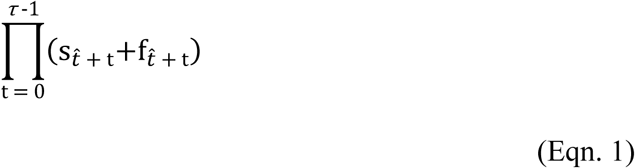

Confidence intervals for these changes were obtained as follows. Parameter values were resampled from the asymptotic multivariate normal distribution specified by the parameter estimates of the Pradel model with lowest AICc. The relative change was calculated for each resample. Among 1000 resamples, the 2.5 and 97.5 percentiles are reported.

To assess estimation bias in the parameters of Pradel models, two approaches were used. 100 datasets per taxon were resampled with replacement, encounter histories constructed and the minimum adequate model fitted to each of these pseudo-datasets. The year-dependent covariates were not resampled. I report estimates for survival probabilities and colonization corrected with bootstrap estimates of bias. The Supplementary Material presents results of further simulations of local survival, colonization, temporary emigration and sampling, which allowed another assessment of estimation bias and of the statistical power to detect decelerating declines and temporary emigration concentrated in a certain period.

## Results

All 105302 and 237542 records from 1946 to 2013 were included in the analysis of *Bombus* and non-*Bombus* genera, respectively. The analysis of the data subsets had (1) 46935 and 207404 records for the analysis of records with specimen in collections and (2) 19763 and 75050 records in grid cells sampled in each of three periods.

## Generalized non-linear models

Prediction intervals of the best maximal models and the models with the most favourable AICc (adequate model) are shown in Figure 2 for the full data and the second subset (first subset results are available in the Supplement). The bootstrap bias assessment shows that the predictions of bias-corrected models remain within the original confidence intervals for *Bombus* but not for the other genera. There, the bootstrap predicts substantial bias. This is most likely due to the fact that I did not add rare species in the bootstrap procedure to compensate for the fact that we can only resample species with records and thus never more species than in the data for that year. Thus, this bootstrap rather shows the relative importance in some years of species with few records in the predicted richness, which is clearly larger for non*-Bombus* genera and for years with fewer records.

**Figure 2.**
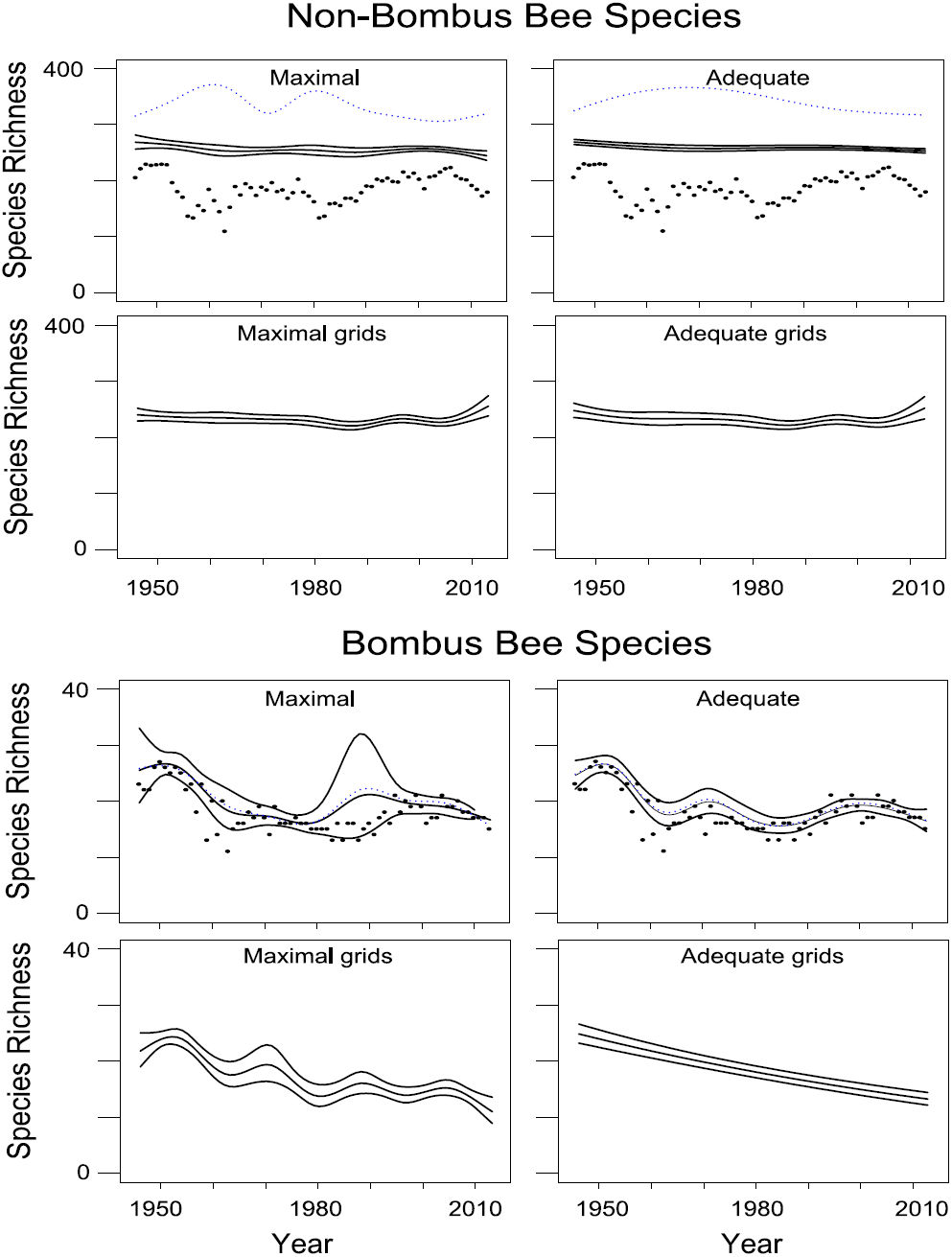
Time patterns of species richness obtained from generalized non-linear modelling (gnlr). Model predictions and 95 % confidence bands of the predicted values are drawn as full lines. Left column: maximum models. Right column: adequate models. For the full datasets, blue dotted lines indicate predictions from a bootstrap-bias-corrected model and raw data points are added for comparison (black). **Top row: all data non-*Bombus* genera**. AICc non-*Bombus* maximal model (*a*(t) 8 d.f. spline of year, *b*(t) 8 d.f. spline of σ): 381.058; AICc non-*Bombus* adequate model (*a*(t) 3 d.f. spline of year, *b*(t) 1 d.f. spline of σ): 365.5. **Second row: data non*-Bombus* genera for grid cells visited in three periods**. AICc non-*Bombus* maximal model (*a*(t) 8 d.f. spline of year, *b*(t) 8 d.f. spline of σ): 390.9; AICc non-Bombus adequate model (*a*(t) 8 d.f. spline of year, *b*(t) 1 d.f. spline of σ): 373.4. **Third row: all data *Bombus***. AICc *Bombus* maximal model (*a*(t) 8 d.f. spline of year, *b*(t) 8 d.f. spline of σ): 308.8; AICc *Bombus* adequate model (*a*(t) 8 d.f. spline of year, *b*(t) constant): 288.0. **Fourth rowh: data *Bombus* for grid cells visited in three periods**. AICc *Bombus* maximal model (*a*(t) 8 d.f. spline of year, *b*(t) 8 d.f. spline of σ): 312.4; AICc *Bombus* adequate model (*a*(t) 1 d.f. spline of year, *b*(t) 3 d.f. spline of σ): 297.0.

A decelerating decrease of species richness occurs when (1) there is a decline in the first half of the interval and (2) any straight line with negative slope drawn within the confidence band in the first half of the study period, runs below the confidence band for the second half. There is no decelerating decrease of species richness in the models for non-*Bombus* genera. The maximal model fitted to the full dataset suggests that it occurs for *Bombus* (Fig. 2). In the maximal models for *Bombus* richness does fluctuate over time, with decreases as good as stalling in the first half of the study period as well, so that it is unclear if any deceleration is real and persists. Comparison with the analyses on the data subset for grids that were repeatedly sampled reveals that a decelerating decline is not found in the adequate model there, and that the richness of non-*Bombus* species did not decline. When models are fitted to the records collected in spatial grid cells that were not repeatedly sampled, the decelerating decline is recovered (Supplement) Thus the conclusion is that this approach is capable of inferring a decelerating decline in species richness, but it is not robustly found. The subset analysis supports that it is generated by spatial sampling heterogeneity over time. Widths of prediction intervals of adequate models suggest that we should not suspect issues of statistical power preventing from detecting decelerations. For non-*Bombus*, species richness in 1946 is estimated from the maximal model as 268 (confidence interval [256, 281]), in 2013 as 244 [236,253]. For *Bombus*, the estimate for 1946 is 26 [20, 33], by 2010 richness is 18 species [17, 18]. Note that the estimated relative loss of richness is larger for *Bombus* than the pooled other bee genera but when relative losses are calculated from confidence intervals limits, they slightly overlap.

## Capture-recapture analysis

Figures three and four show the results of fitting time-dependent (left, no mixtures) or regression Pradel models with mixtures (right) to the data. Reducing the dataset to a shorter interval (1957–2002) showed in time-dependent models that colonization was again increased at the new beginning of the shorter interval and that survival decreased near the new end, revealing that the similar estimated survival and colonization patterns near the start and the end of the entire study period are to be seen as biased. In both groups of species, models with detection probabilities that are mixtures of several components were preferred (Table 1, Figures 3 and 4). The time-dependent models show negligible colonization and years with reduced survival. In the case of *Bombus*, the absence of a year with decreased survival between 1985 and 2000 makes the species richness decline decelerate in the time-dependent model: there just isn’t a downward step in that period (Figure 4, left column). However, the number of such years is only 4 between 1955 and 2003, therefore a waiting time of this length shouldn’t be taken as conclusive evidence of a changed process. Table 1 gives AICc values of the adequate models and a set of other models that are useful for comparison. In these tables, “*t*” indicates time categorical effects fitted, “*T*” a linear effect in the linear predictor, which was linked to the data using log (colonization) or logit (local survival) link functions. The number of mixture components in the detection model is given. In the last two columns of the tables, estimates of the linear effect of *T* are given when fitted. At least one estimate for local survival or colonization needs to be significantly positive for a deceleration in the decline of species richness to be possible. This does not occur, also not in analysis of the different data subsets (Table S2). However, this is in agreement with simulation results (Supplementary Material) that indicate that there is a bias towards estimating negative trends in local survival. When time-dependent and regression models are compared (no mixtures, Figs. 3 and 4, second rows), there are a limited number of years where detection probabilities differ significantly between these models, and only in the non*-Bombus* genera. These years in between 1955 and 1965 and after 2000 could be years where a fraction of species temporarily emigrated. However, the number of such years is limited and their distribution over time such that a decelerating decline cannot be inferred from them.

**Table 1.**
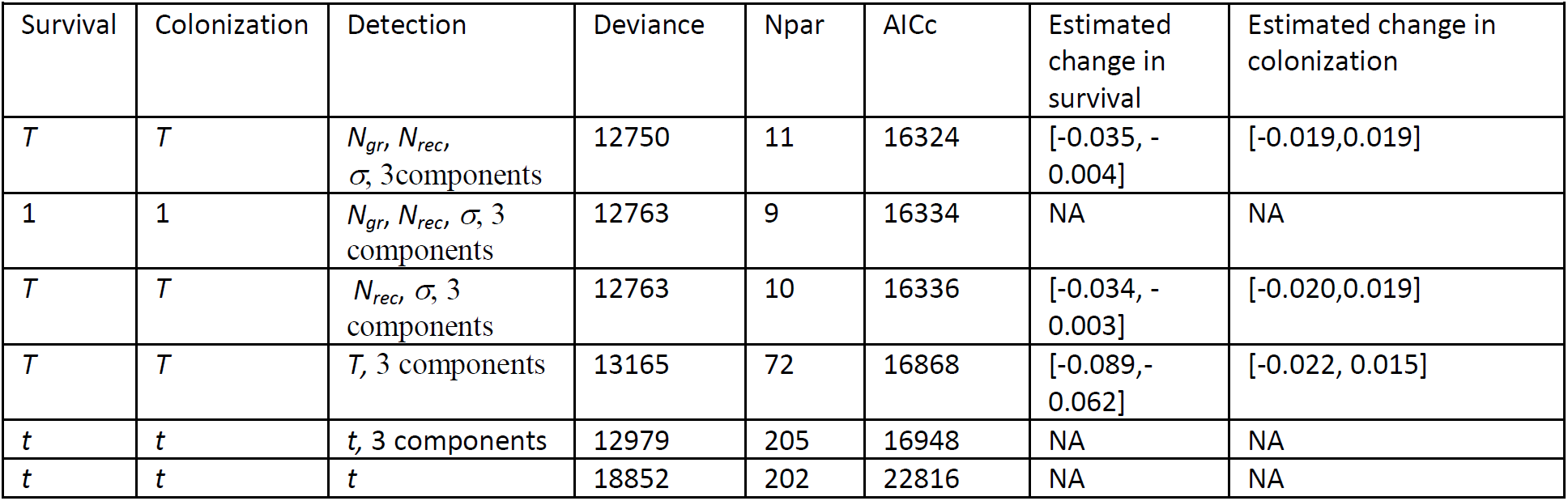
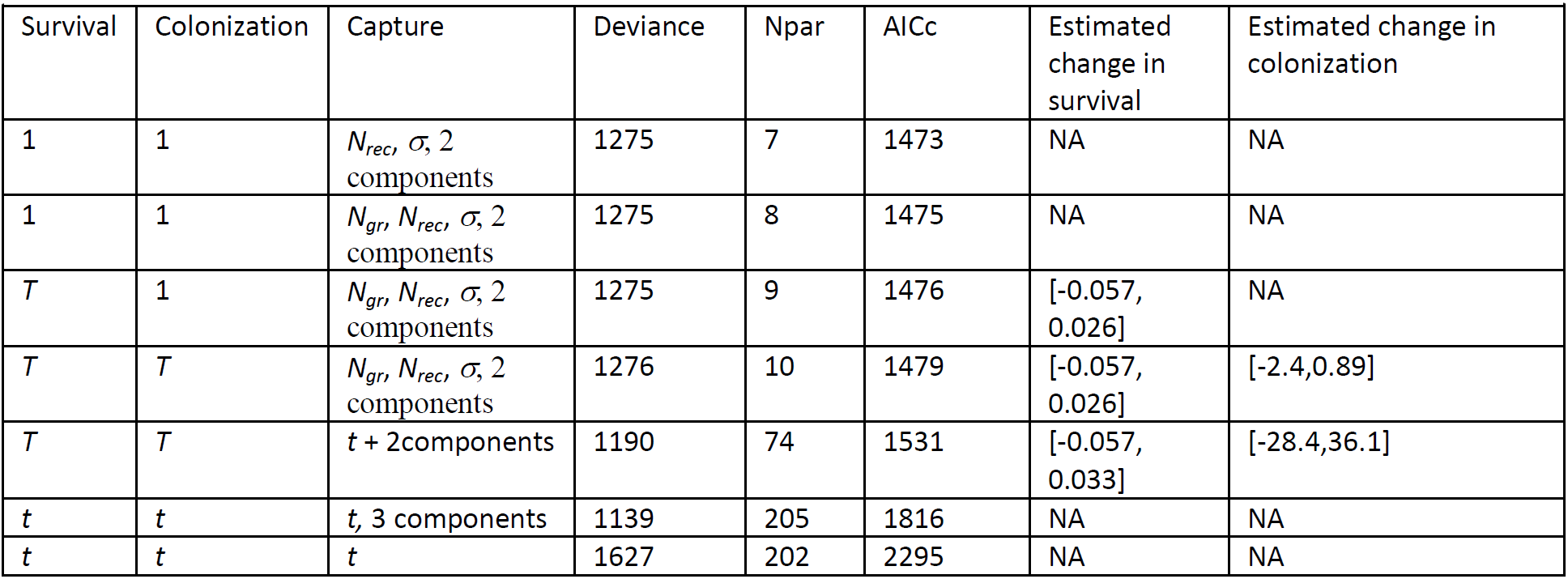
Pradel models fitted to the data. The first three colums list the covariates in the models for local survival, colonization and detection, the last two columns confidence intervals for the time trends in survival and colonization, when estimated. *T* refers to a linear effect of year, *t* to categorical effects of year, “ *1”* indicates a model without explanatory variables. The numbers of components in the mixture distributions for detection probabilities are given. Abbreviations for the year-dependent variables are *σ* for the heterogeneity parameter of the poisson-log normal distribution, *N_rec_* is the total number of records per year, *N_grid_* the number of 10 by 10 km grid cells sampled.

| Table 1a. Non-*Bombus* Bees - All data 1946-2013 |  |  |  |  |  |  |  |
| --- | --- | --- | --- | --- | --- | --- | --- |
| Survival | Colonization | Detection | Deviance | Npar | AICc | Estimated change in survival | Estimated change in colonization |
| $T$ | $T$ | $N_{gr}, N_{rec}, \sigma, 3$ components | 12750 | 11 | 16324 | [-0.035, -0.004] | [-0.019, 0.019] |
| 1 | 1 | $N_{gr}, N_{rec}, \sigma, 3$ components | 12763 | 9 | 16334 | NA | NA |
| $T$ | $T$ | $N_{rec}, \sigma, 3$ components | 12763 | 10 | 16336 | [-0.034, -0.003] | [-0.020, 0.019] |
| $T$ | $T$ | $T, 3$ components | 13165 | 72 | 16868 | [-0.089, -0.062] | [-0.022, 0.015] |
| $t$ | $t$ | $t, 3$ components | 12979 | 205 | 16948 | NA | NA |
| $t$ | $t$ | $t$ | 18852 | 202 | 22816 | NA | NA |
| Table 1b. *Bombus* Bumblebees - All data 1946-2013 |  |  |  |  |  |  |  |
| Survival | Colonization | Capture | Deviance | Npar | AICc | Estimated change in survival | Estimated change in colonization |
| 1 | 1 | $N_{rec}, \sigma, 2$ components | 1275 | 7 | 1473 | NA | NA |
| 1 | 1 | $N_{gr}, N_{rec}, \sigma, 2$ components | 1275 | 8 | 1475 | NA | NA |
| $T$ | 1 | $N_{gr}, N_{rec}, \sigma, 2$ components | 1275 | 9 | 1476 | [-0.057, 0.026] | NA |
| $T$ | $T$ | $N_{gr}, N_{rec}, \sigma, 2$ components | 1276 | 10 | 1479 | [-0.057, 0.026] | [-2.4, 0.89] |
| $T$ | $T$ | $t + 2$ components | 1190 | 74 | 1531 | [-0.057, 0.033] | [-28.4, 36.1] |
| $t$ | $t$ | $t, 3$ components | 1139 | 205 | 1816 | NA | NA |
| $t$ | $t$ | $t$ | 1627 | 202 | 2295 | NA | NA |

**Figure 3.**
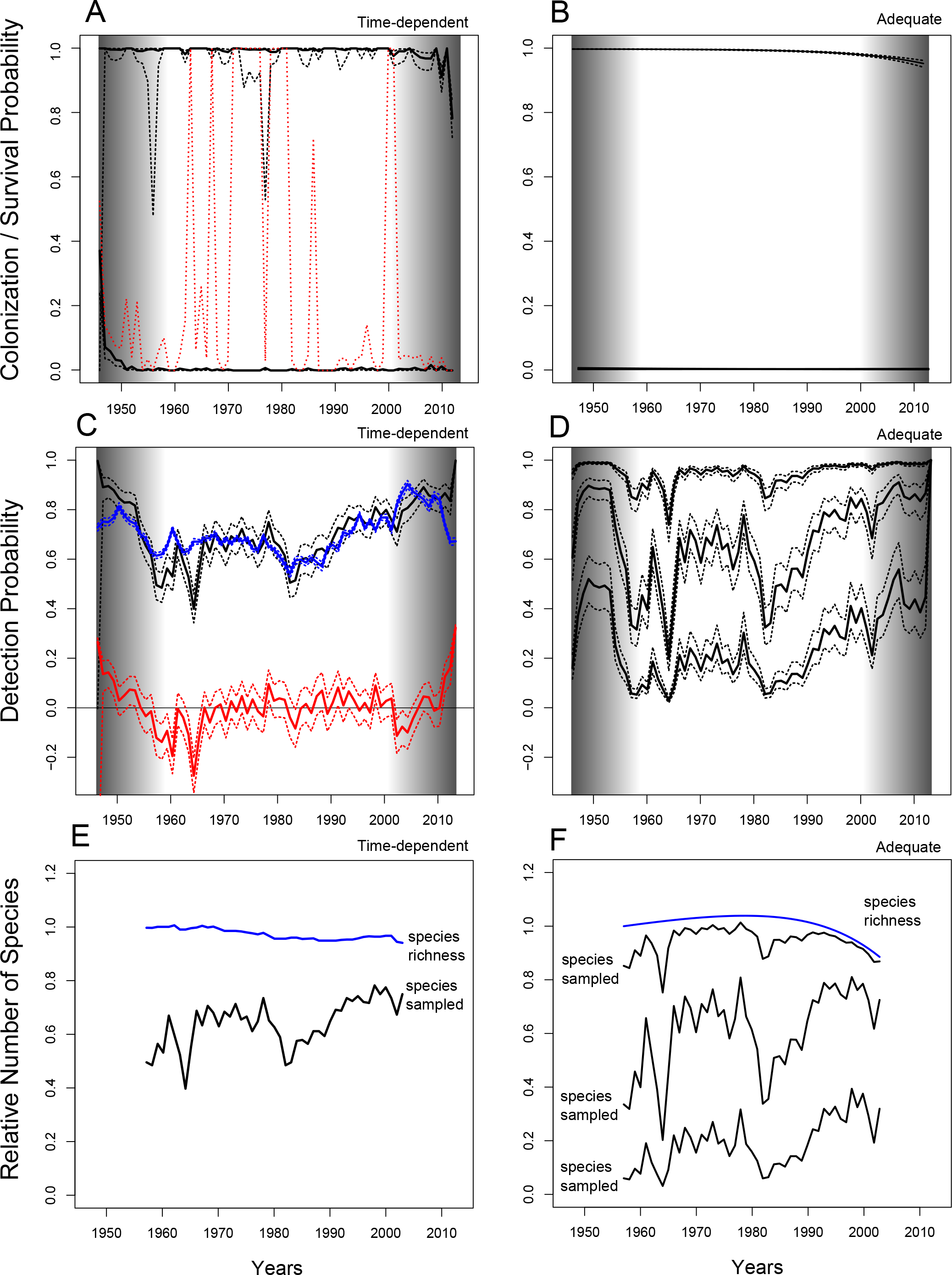
Results obtained from fitting Pradel models to species from non-*Bombus* wild bee genera. Left column: Parameter estimates and predictions of the fully time-dependent model without fitting mixtures; right column: the adequate model with a mixture for detection probability (Table 1). Top row: Local survival (upper full curve) and colonization probabilities (lower full curve) per year, which are close to values one and zero, respectively. Confidence intervals are indicated as dotted lines, in red for colonization probabilities. Middle row: Detection probabilities. In the left panel the predictions of a model with categorical time effects (black) and a model with logit regressions of the three year-specific explanatory variables (blue) are plotted with 95% confidence intervals. The differences between both are plotted in red and can be informative on the occurrence of temporary emigration. Right panel: For the adequate model, detection probabilities vary between species, and the curve plotted is for each of the three components in the species mixture. Bottom row: Trends in relative species richness over years, where 1955 is used as a reference and given value 1 (blue: species richness, black: for comparison, predicted relative number of species sampled). In the bottom right panel, the predicted relative number of species sampled is plotted for the mixture components as well. Grey bands: biased parameter estimates are expected for these years on the basis of a similar analysis over a shortened time interval.

**Figure 4:**
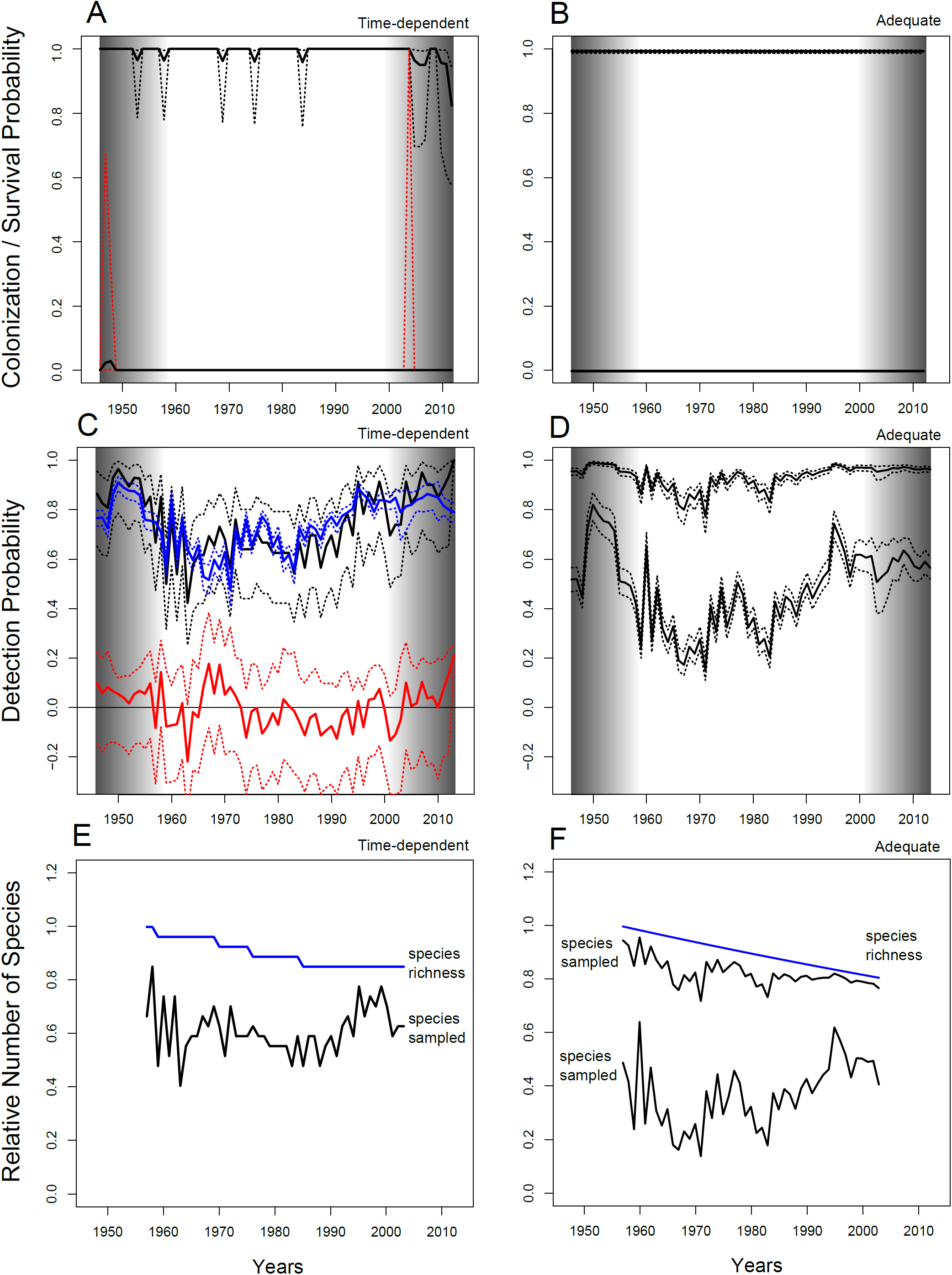
Results obtained from Pradel models fitted to species in the *Bombus* bumblebee genus. Order and content of all panels are as for Figure three. For *Bombus*, detection probability is a mixture of two components (groups of species). Additional information on the adequate model can be found in Table one.

Correcting parameter estimates of linear time trends in survival and colonization with a bootstrap estimate of bias gave the following results. For non-*Bombus* the bias-corrected estimate for the change in survival probability (on the logit scale) equals -0.014, slightly larger than the uncorrected estimate. The bias-corrected estimate of the time trend parameter for colonization is -7.52, of little relevance as the estimate basically implies the absence of colonization. For *Bombus* the bias-corrected survival trend is -0.017, the trend for colonization -0.91. The bootstrap estimates of bias are not consistently in agreement with the results of the separate simulations and therefore seem of limited value. Based on parameter estimates of the models with lowest AICc, in 2017 a predicted 9.6% (confidence interval [0, 19%] and 29% [15, 51] of species of non-*Bombus* wild bees and *Bombus* bumblebees, respectively, have disappeared since 1946. Simulations (Supplement) indicate that these decreases are probably underestimating the real decreases, but a comparison with the results of generalized linear modelling suggests that the underestimation might be limited.

## Discussion

A dataset with records and observations of wild bee species in the Netherlands was analysed using different approaches to investigate trends in species richness for *Bombus* and non-*Bombus* genera across almost 70 years. Neither of these approaches detected a decelerating trend with sufficient consistency to reject the hypothesis of a constant decline or no decline at all. Generalized non-linear modelling showed potential estimation bias for non-*Bombus* genera, which followed directly from limitations on the bootstrap procedure, as absent species can not be resampled. I therefore take this bias rather as an indication of the importance of species with few records in the results. Simulations indicated that the modelling of species richness changes via encounter histories suffered from estimation bias of parameters representing time trends, making it difficult to detect decelerating declines using that method altogether. For generalized linear modelling, a similar time trend was estimated in different subsets, and no spurious decelerating declines were caused by model selection bias.

The fraction of species richness lost since 1946 or the species richnesses at the end of the study period could be predicted for either approach. From the analyses, the conclusion is that species richnesses have declined since 1946 for the *Bombus* genus. For the non-*Bombus* genera analysed using encounter histories, the 95% confidence interval touches zero but does estimate a small decrease. Similarly the generalized linear models estimate a small or no decline.

## Wild bee species richness trends

I found that species richness in the *Bombus* genus is substantially reduced, and the decrease estimated is larger than in a study using rarefied richness on Swedish data (Bommarco et al. 2011). On the other hand, the estimated overall decrease for the other genera seems smaller than in an analysis of bee species richness in the UK with relative decreases between 10 and 30% in most study sites, when comparing periods separated by 33 years (Senapathi et al. 2015). However that study used the same methods as Carvalheiro et al. (2013) and no confidence intervals for the changes were given. Again for the UK, Ollerton et al. (2014) analysed bee and flower-visiting wasp species richness and noted extinctions before 1960, which they attributed to agricultural intensification. These authors noted that their results for the most recent decades contradict Carvalheiro et al. (2013), as extinctions might be increasing again. In the methods used here, the most adequate models find continuing declines, potentially with fluctuations over time.

Risks of species richness loss seem taxon-specific (*Bombus* vs. non-*Bombus*), and thus call for trend analysis for smaller taxonomic groups in general. However that requires sufficient data for each of them. The stronger decrease suggested for *Bombus* is surprising if it is correct, given that bumblebee densities and presences are assumed not to be determined by very local landschape characteristic and can benefit from urbanisation (Carré et al. 2009; Kennedy et al. 2013; Senapathi et al. 2016). However, bumblebees might show an elevated susceptibility to rapid climate change (Kerr et al. 2015).

Combining the results on Dutch wild bees and bumblebees with the reassessment of Van Dooren (2016), the conclusion must be that there are no decelerations in the declines of species richness in pollinators in North-West Europe. There is thus no reason to be satisfied with current biodiversity conservation efforts, as there is no or insufficient evidence that they have been effective. It has been stated elsewhere that the most recent EU CAP Agricultural Reform fails on managing biodiversity adequately (Pe’er et al 2014). Even while pollination services are provided by the abundance of the most common species (Kleijn et al. 2015), we should not forgo protecting the rarer ones. Inferring arrested biodiversity declines as was done by Carvalheiro et al. (2013) appears dangerous.

## Methodogical developments

Boyd (2013) has proposed that audited standards robust to variations in assessor competence should be available and used in biodiversity research and data collection. For historical data, it is too late for that. Retrospective sampling standardization is impossible. With data of the type analysed here, we therefore need to resort to detailed statistical modelling, with conservative inference to avoid new erroneous conclusions. The situation could have been as bad as O’Hara (2005) suggested, namely that we often rather analyse properties of estimators than richness patterns themselves. This is confirmed for the estimation of time trends in local survival using Pradel models, where estimation bias leads to wrong inference and masks differences between datasets. The relatively simple generalized non-linear model thus becomes the most dependable tool to infer species richness trends for now. It does not need to assume constant temporary emigration, and the manner in which the sampling process is modelled comes with limited assumptions. However, the method needs manual work, as model fitting is tedious. Convergence is not guaranteed and results need to be inspected with care.

The results of each analysis presented here urge further methodological developments. Chao et al. (2014) proposed a bootstrap method to estimate variances of extrapolated richness, where the fraction of rare missing species is first estimated and then added to an imputed dataset from which resampling occurs. I have used bootstrap methods differently, to estimate magnitudes of estimation bias in the different types of analysis. The idea there is that the estimates based on the resamples differ from the estimate on the actual data in the same way as the estimate differs from the true value (Davison & Hinkley 1997). The approach found estimation bias in non-linear regressions for non-*Bombus*. It should be further verified whether bias was adequately estimated using this bootstrap or just a side effect of the importance of species with few records in a non-random sample. For the capture-recapture analysis, bootstrap bias estimates did not very well align with expectations from simulations.

Issues with models for encounter histories will probably be alleviated when robust design models (Pollock 1982) can be fitted to the data. If records had been collected throughout the study period independently by observations or via specimens deposited in collections, then these two sampling methods could have been used to fit a robust design. The pattern of estimation bias we found might also be specific to this dataset.

Heterogeneity in species detection probabilities can be expected in all methods addressing species richness (e.g. Boulinier et al. 1998) and it was incorporated into the models by means of mixtures. Note that models for open populations were used in the analysis of species encounter histories. Models exist to infer species richness in open occupancy models (Nichols et al. 1998; Yamaura et al. 2011), but these often depend on the assumption of individual random sampling which is untenable here. Statistical modelling of how collectors vary in their sampling efforts over time clearly deserves more study and wider application, given that more and more data of the kind analysed here are mined and as people are encouraged to collect citizen-science data (Potts et al. 2016b). Also studies on abundance trends in different taxa (e.g. Inger et al. 2014) could benefit from multi-model inference and assessments of bias-variance tradeoffs and sampling heterogeneity.

To conclude, I want to point out a potential example of how non-random data collection might be generated. Note that a recent Dutch reference work on wild bees (Peeters et al. 2012) explicitly points to recent records of the wild bee species *Andrena coitana* in Germany and the habitat type in the Netherlands where the species might be seen again after a long period without records. Recently, the rediscovery of *Andrena coitana* was reported (Nieuwenhuijsen 2016). We cannot exclude the possibility that a new reference work or other species exposure motivates an increased effort to collect particular species. This can thwart any effort to achieve random sampling, so essential to most of our inference methods, by replacing it with recording syndromes (Isaac & Pocock 2015).

## Acknowledgements.

I thank Menno Reemer for providing the EIS data. Roger Pradel gave useful comments on a previous version.

## Supplementary Material

Complete and self-explanatory R scripts of the analysis and data frames with variables are available from the author as a compressed .zip file. While the main text discusses two approaches to modelling species richness change, here I present four. The first is mainly there to illustrate that counts are the natural kind of random variables to represent richnesses. The third was an attempt to integrate estimates of species richness obtained using independent software into a regression framework.

### Modelling species number accounting for sample coverage

Chao and Jost (2012) have proposed to compare species numbers at fixed sample coverage. Using species number data, one can construct a simple linear model that includes years and sample coverage as explanatory variables, and which predicts species richness at sample coverage equal to one.

For species numbers “sp”, years “Years” and sample coverage “Coverage”, and assuming a simple quadratic model for the year effects the model is

~~~
model<-glm(Sp~Years+Years^^^2+I(1-Coverage),family=poisson)
# predicted richnesses at sample coverage = 1
predicted_richness<-
predict(model,newdata=list(coverage=rep(1,69)),type=“response”,se.fit=T)
~~~

I did not pursue this generalized linear modelling approach further, for example by modelling standard deviations of species number as below. It is unlikely that the data in this paper were obtained using random sampling. Estimates of sampling coverage might therefore be inadequate and their precision difficult to assess. The correction for coverage is linear, therefore the range of coverages in the data should be limited to make this approximation valid. This method might perform well for randomly sampled data and different subsets per year with slightly different coverages so that there is coverage variation independent of years.

### Generalized non-linear regression

This approach and some of its characteristics are detailed in the main text. None of the estimation methods proposed by Raaijmakers (1987) nor the non-linear Least Squares used by O’Hara (2005) were applied. Here I paste the code for one of the most elaborate models fitted using gnlr (Lindsey 1997), including the initial values for all parameters used in the ML optimization. In the short script below, “Sp” is the number of species, “spl8” is a natural cubic spline of eight d.f. of the year variable, “rec” is the number of records per year. “SD” is the predicted unconditional standard deviation of the number of species at the actual number of records, calculated following Colwell et al. (2012).

~~~
gnlr(sp,mu=~exp(a+a1*spl8[,1]+a2*spl8[,2]+a3*spl8[,3]+a4*spl8[,4]+a5*spl8[,5]+a6*spl8[,
6]+a7*spl8[,7]+a8*spl8[,8])*rec/(rec+exp(b+b1*spl8[,1]+b2*spl8[,2]+b3*spl8[,3]+b4*spl8,
4]+b5*spl8[,5]+b6*spl8[,6]+b7*spl8[,7]+b8*spl8[,8])),
pmu=c(1,0,0,0,0,0,0,0,0,1,0,0,0,0,0,0,0,0),
shape=~log(exp(v0)+SD^2),pshape=list(v0=-2))
~~~

Full scripts with all models and the model selection procedures can be provided as text files.

### Smooth regressions of species richness estimates

In this third type of analysis, I used species richness estimates in smooth regression models with several covariates. As estimates of species richness, I used the “best” estimator selected among five different parametric models using a detailed heuristic (Bunge et al. 2012). This approach thus does not have a clear part of the model representing the observation process, except for the fact that some covariates related to it are used in smooth regressions. Program CatchAll (Bunge et al. 2012) was used to obtain these estimates of species richness per year, with an estimated standard deviation. Time patterns in log annual species richness estimates were then modelled using smooth functions of the following explanatory variables: the year where the samples or records were from, the number of 10×10 km grid cells with records per year, and the estimated variance parameter a of the Poisson lognormal distribution fitted to numbers of records per species per year for the taxonomic group analysed. These smooth regressions of log-transformed species richness were fitted using the gamlss() function for R (Stasinopoulos and Rigby 2007). I modelled heterogeneous errors in log species richness. I fitted as maximal models cubic splines of 9 d.f. for the year variable, and of 4 d.f. for the other explanatory variables (note that d.f. in gamlss() output are given as on top of the linear model). First the model was simplified by reducing the d.f. of the cubic spline for the year variable, subsequently by doing that for the other variables. Models were compared using AICc as for the models in the main text, until a minimum adequate model with lowest AICc was identified.

For the estimation bias assessments in this analysis, 100 datasets were re-sampled per year and per taxonomic group and the best species richness estimate determined for each resample. The average estimated species richness per year across re-samples was then used in a bias correction of the original species richness estimate of that year. The gamlss() models were refitted to the bias-corrected data with the standard errors of predicted log species richness also adjusted to the additional calculations. In figures, the smooth effects of the covariates on predicted log species richnesses are transformed back to the original scale, such that they center around value one.

In the maximal models, species richness again showed a trend with fluctuations across years (Fig. S1, second and fourth row). For the non-*Bombus* data (Table S1, Fig. S1), there are no declines detected and even an increase in species richness is inferred for one adequate model (Fig. S1, Right column). In *Bombus*, a decline is consistently found, and it decelerates in one data subset (Fig. S1, Middle column) which can be interpreted as following from model selection bias. An analysis with bootstrap-bias-corrected species richness estimates did not suggest substantial estimation bias for any of the models nor taxa.

I decided not to present this method in the main text as it has no modelling of the sampling process, it is only the “true” species richness that is predicted. The method might perform adequately when parametric distributions of species abundances are well known and stable across the study period, when there is no need to predict species richness from partial data, i.e., when samples have perfect coverage.

A maximal model fitted in R using the gamlss() function was written at follows:

~~~
gamBPnb8<-gamlss(formula=log(pred)~cs(year,df=8)+cs(poilogsigma)+cs(Ngrids),
sigma.formula=~offset(exp(predse/pred)))
~~~

Here, “pred” is the predicted species richness, “year” is the calendar year, “poilogsigma” is the sigma parameter, “Ngrids” the number of 10×10 km grid cells with samples in each year and “predse” is the estimated standard error of the predicted species richness.

To obtain a bootstrap estimate of bias, the bias-corrected species richness “predcorr” was fitted with the following modified expression for its standard deviation “predcorrse”, where “bootav” stands for the mean of the bootstrap estimates of species richness, “bootse” for the standard deviation of the boostrap estimates of species richness, and “nsamples” for the number of bootstrap samples drawn.

~~~
predcorr<-2*pred-bootav
predcorrse<-sqrt(4*predseA2+bootseA2/nsamples)
gamBPnb8<-gamlss(formula=log(predcorr)~cs(year,df=8)+cs(poilogsigma)+cs(Ngrids),
sigma.formula=~offset(exp(predsecorr/predcorr)))
~~~

**Figure S1.**
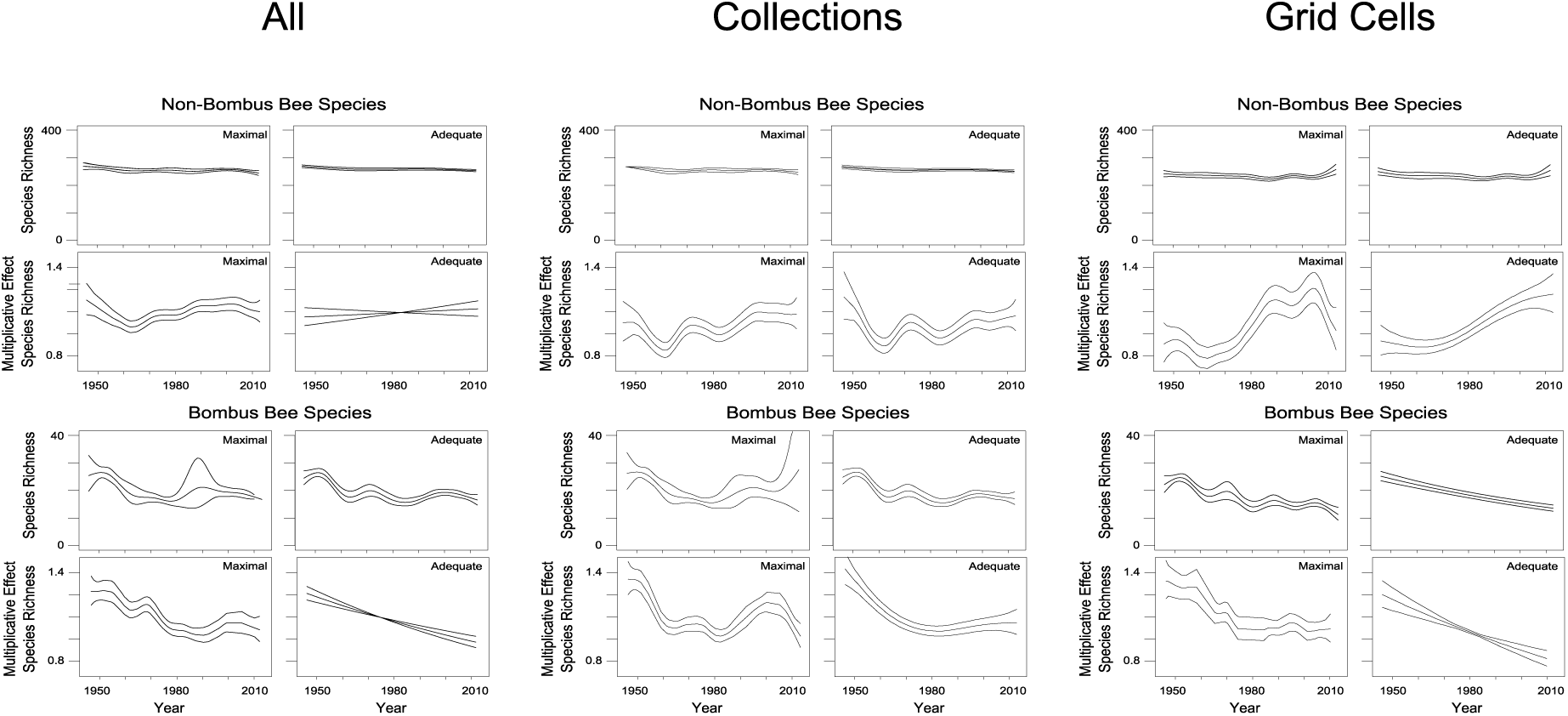
Results when modelling different subsets separately. Results of generalized non-linear models and smooth regressions. Analysis of the full datasets in the left column. Non-*Bombus* and *Bombus* records for which specimens were deposited in museum collections (middle), for grid cells sampled in each of the three time periods (right). Estimates of species richness (gnlr) and of the relative change in species richness across years (gamlss). Selected models are given in Table S1 below.

**Table S1.** Adequate models obtained for the gnlr and gamlss analyses of data subsets. For gnlr, the explanatory variable and the degrees of freedom of the splines used to model *a*() and *b*() are given. For gamlss the spline degrees of freedom of the variables in the selected regression models.

| non-*Bombus* | All | Collection | Grids |
| --- | --- | --- | --- |
| Gnlr | $a(\text{year}, \text{d.f.} = 3)$<br>$b(\sigma, \text{d.f.} = 1)$ | $a(\text{year}, \text{d.f.} = 3)$<br>$b(\sigma, \text{d.f.} = 1)$ | $a(\text{year}, \text{d.f.} = 8)$<br>$b(\sigma, \text{d.f.} = 1)$ |
| Gamlss | Year d.f. = 1<br>$N_{grid}$ d.f. = 4<br>$\sigma$ d.f. = 2 | Year d.f. = 8<br>$N_{grid}$ d.f. = 1 | Year d.f. = 3<br>$N_{grid}$ d.f. = 3<br>$\sigma$ d.f. = 2 |
| *Bombus* |  |  |  |
| Gnlr | $a(\text{year}, \text{d.f.} = 8)$<br>$b(\text{constant})$ | $a(\text{year}, \text{d.f.} = 8)$<br>$b(\text{constant})$ | $a(\text{year}, \text{d.f.} = 1)$<br>$b(\sigma, \text{d.f.} = 3)$ |
| Gamlss | Year d.f. = 1<br>$\sigma$ d.f. = 1 | Year d.f. = 3<br>$N_{grid}$ d.f. = 4 | Year d.f. = 1<br>$\sigma$ d.f. = 1 |

**Table S2.**
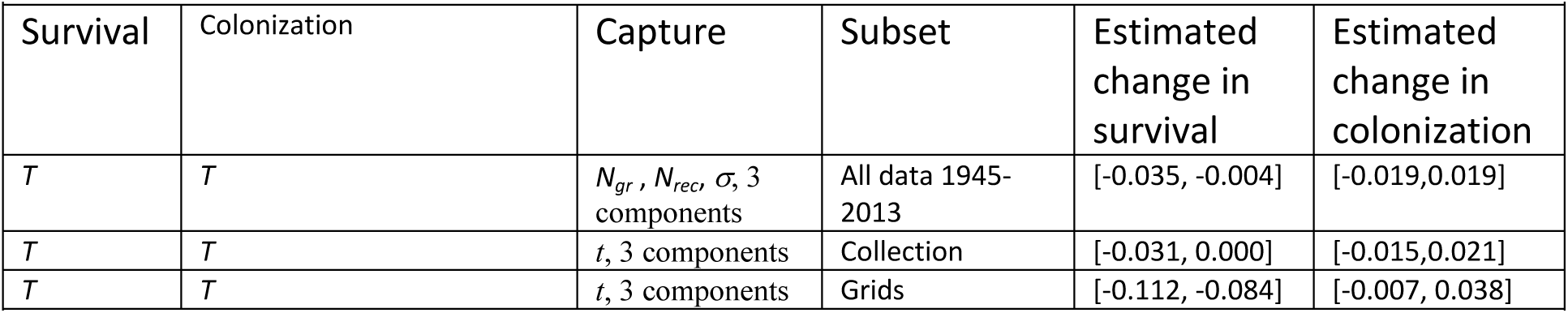
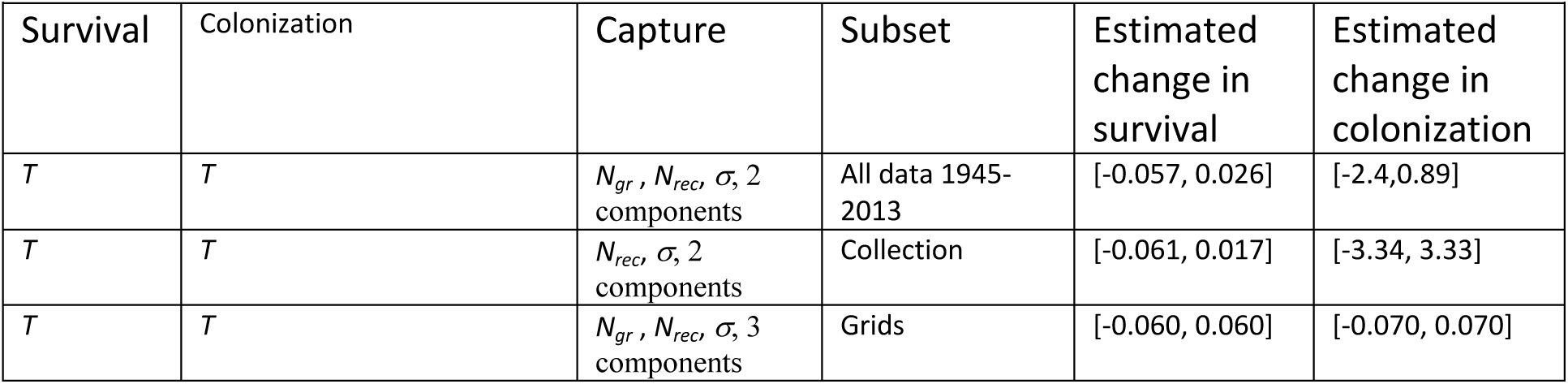
Pradel models fitted to data subsets. Minimum adequate models are given among the models with estimates of time trends of colonization and survival. Abbreviations of explanatory variables are as in Table 1.

| Table S2a. Non-*Bombus* Bees – Time trends for each subset analysis |  |  |  |  |  |
| --- | --- | --- | --- | --- | --- |
| Survival | Colonization | Capture | Subset | Estimated change in survival | Estimated change in colonization |
| $T$ | $T$ | $N_{gr}, N_{rec}, \sigma, 3$ components | All data 1945-2013 | [-0.035, -0.004] | [-0.019, 0.019] |
| $T$ | $T$ | $t, 3$ components | Collection | [-0.031, 0.000] | [-0.015, 0.021] |
| $T$ | $T$ | $t, 3$ components | Grids | [-0.112, -0.084] | [-0.007, 0.038] |
| Table S2b. *Bombus* Bumblebees – Time trends for each subset analysis |  |  |  |  |  |
| Survival | Colonization | Capture | Subset | Estimated change in survival | Estimated change in colonization |
| $T$ | $T$ | $N_{gr}, N_{rec}, \sigma, 2$ components | All data 1945-2013 | [-0.057, 0.026] | [-2.4, 0.89] |
| $T$ | $T$ | $N_{rec}, \sigma, 2$ components | Collection | [-0.061, 0.017] | [-3.34, 3.33] |
| $T$ | $T$ | $N_{gr}, N_{rec}, \sigma, 3$ components | Grids | [-0.060, 0.060] | [-0.070, 0.070] |

### Generalized linear models fitted to grid cells that were not repeatedly sampled in different time periods

When generalized linear models are used to analyse the dataset restricted to grid cells that were sampled early, middle and late in the study period, a decelerating decline is not retained in the minimum adequate model. To investigate whether the decelerating decline in the full dataset is due to grid cells that were not repeatedly sampled across the study period, these are analysed separately as well. Here I report on the analysis of the dataset consisting of records from grids that were not in the subset presented in Figure two (rows two and four). These data consist of 162492 records of non-*Bombus* bees, and 85539 records of *Bombus* bumblebees.

Among the maximal models for the non-*Bombus* genera, the AICc for the model (*a*(t) 8 d.f. spline of year, *b*(t) 8 d.f. spline of σ) was lowest, AICc = 393.9. The minimum adequate model was (*a*(t) 6 d.f. spline of year, *b*(t) 3 d.f. spline of σ) with AICc equal to 382.4.

Among the maximal models for the *Bombus* bumblebees, the AICc for the model (*a*(t) 8 d.f. spline of year, *b*(t) 8 d.f. spline of σ) was lowest, AICc = 302.9. The minimum adequate model was (*a*(t) 8 d.f. spline of year, *b*(t) 0 d.f. spline of σ) with AICc equal to 281.5.

Figure S2 shows that decelerating declines are retained in the adequate model for *Bombus*.

**Figure S2.**
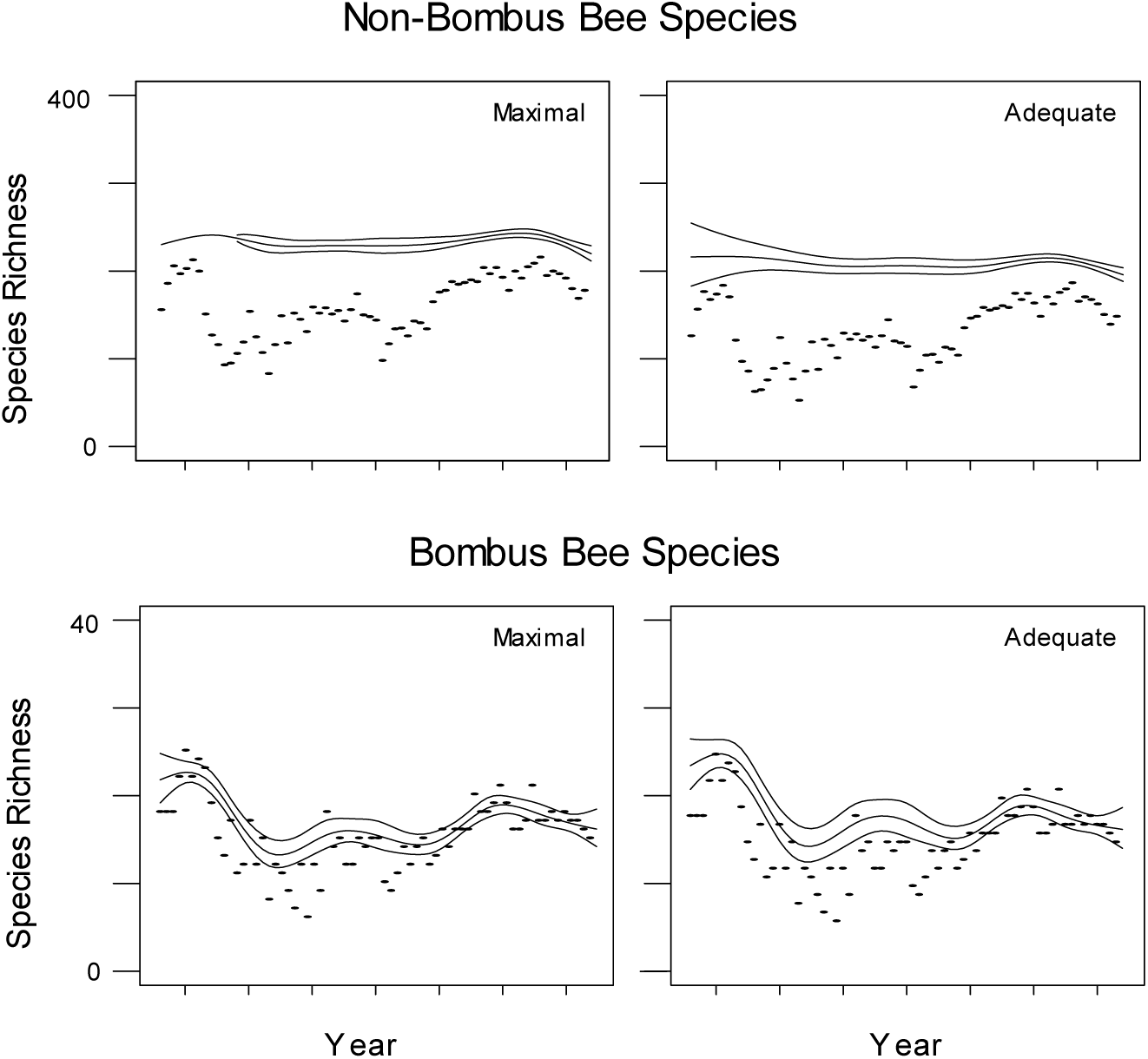
Time patterns of species richness obtained from generalized non-linear modelling (gnlr) on the subset of the data restricted to grids that were not repeatedly sampled as explained in the main text. Model predictions and 95 % confidence bands of the predicted values are drawn as full lines. Raw species count values are added as points. Left column: maximal models. Right column: adequate models. Maximal and adequate models and their AICc are given in the text above.Simulations of species change and estimation

Changes in bee species richness were simulated using the relative numbers of individual records per species in the data as proxy for the initial relative abundances. All simulations started with 324 non-*Bombus* or 28 *Bombus* species. Only species local survival was modelled, and no colonization. Simulation code is provided as a text file. Simulated assemblage time series were randomly sampled to match the numbers of records per year in the actual data and to produce time series of simulated records. Different effects on local survival were considered 1) different time trends in species survival probabilities 2) effects of relative abundance on the probability that a species survives 3) different average survival probabilities (intercepts) at the start of the time series.

To simulate survival, we drew per year and species a random binary variable with average probability of success (survival) of

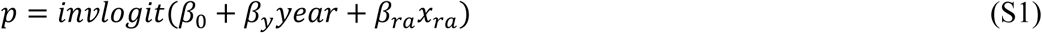

With *year* for the year regression effect and with *x_ra_* for the relative abundance of the species in that year. Parameter values for *β*_0_ were (4, 5, 6, 7), for *β*_1_ (-0.02, 0, 0.02, 0.05), for *β*_2_ (0, 0.0005, 0.0010). Per combination of parameter values, three replicates were simulated and simulations were repeated for *Bombus* and non-*Bombus* genera separately, with similar results. To each simulated dataset, different Pradel models were fitted. All estimated a constant colonization probability. Local survival was either constant, with a regression of the year effect, with an effect of the total number of records per species, with additive effects of both. Detection probabilities were fitted with a regression of the year effect, with effects of *σ* and the number of observations, with the last two effects and the total number of records. A model with constant survival and colonization and a regression for detection probability with *σ* and the number of observations was usually preferred. Only results for the non-*Bombus* genera are shown below.

The main results are the following, for the models that do estimate year effects. First, time trends in disappearance are underestimated across models, with stronger underestimation of positive trends in the probability to survive and stronger underestimation when the local survival probability per year is large (Fig S3). Second, immigration was overestimated. The intercept for colonization per year was estimated to be 0.0038 on average (s.d. 0.0018), while we did not simulate colonization at all. Third, when species richness change across the entire simulation period (50 years) is estimated, including recruitment in the calculation leads to underestimation of the decrease in richness across the period, setting recruitment to zero leads to overestimation of the decrease (Fig S4). Fourth, the intercept estimate for local survival appeared relatively unbiased.

**Fig S3.**
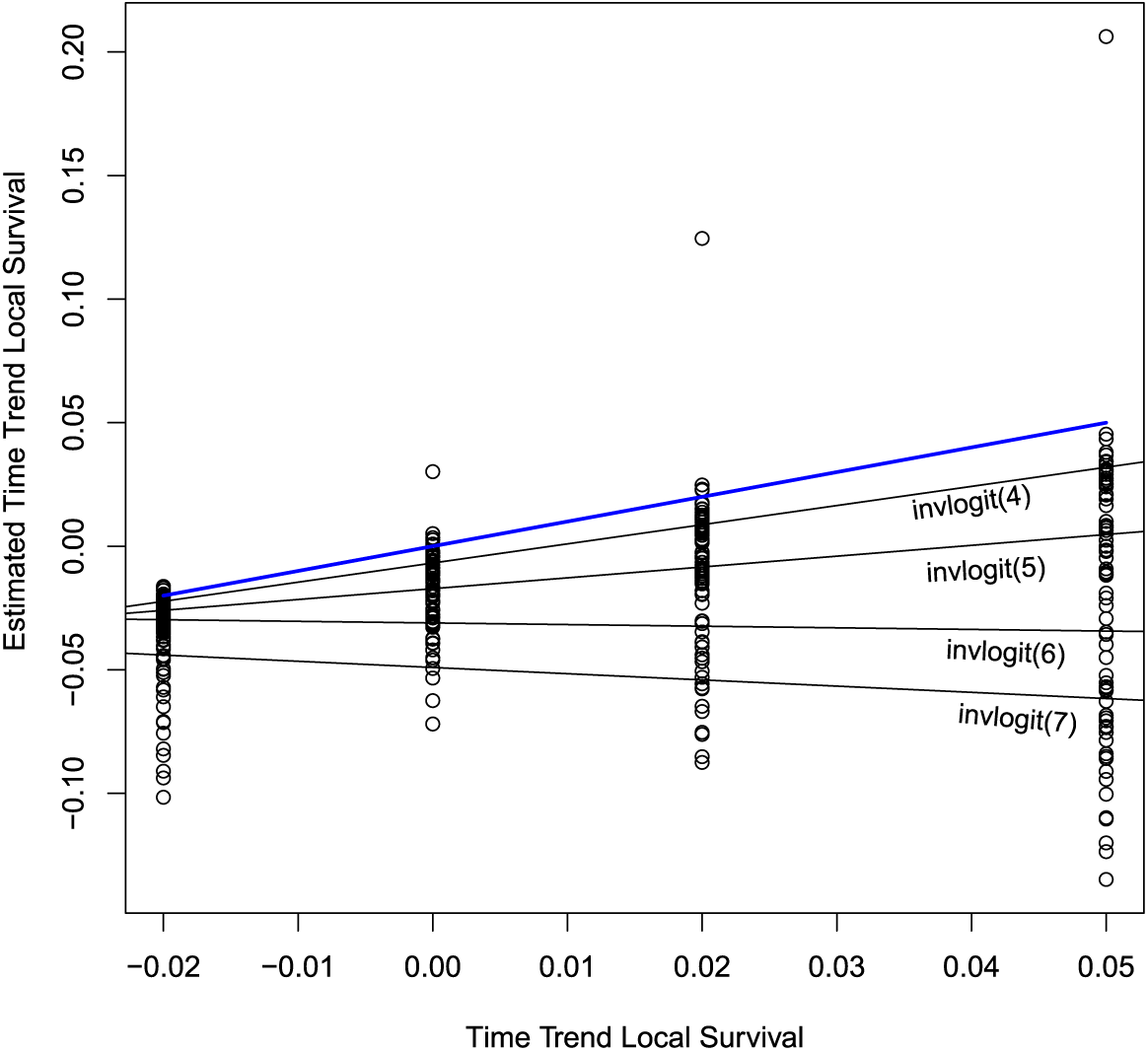
Estimates of the time trend in local survival, across all simulations without effects of relative abundance on survival. All estimates of models fitted with year regression effects are shown, thus including estimates for six different models fitted to each simulated dataset. Different regression models were fitted to the estimates per simulated intercept for local survival, showing the effect on underestimation. For reference, the true value in the simulations of the time trend is indicated by a blue line.

**Fig S4.**
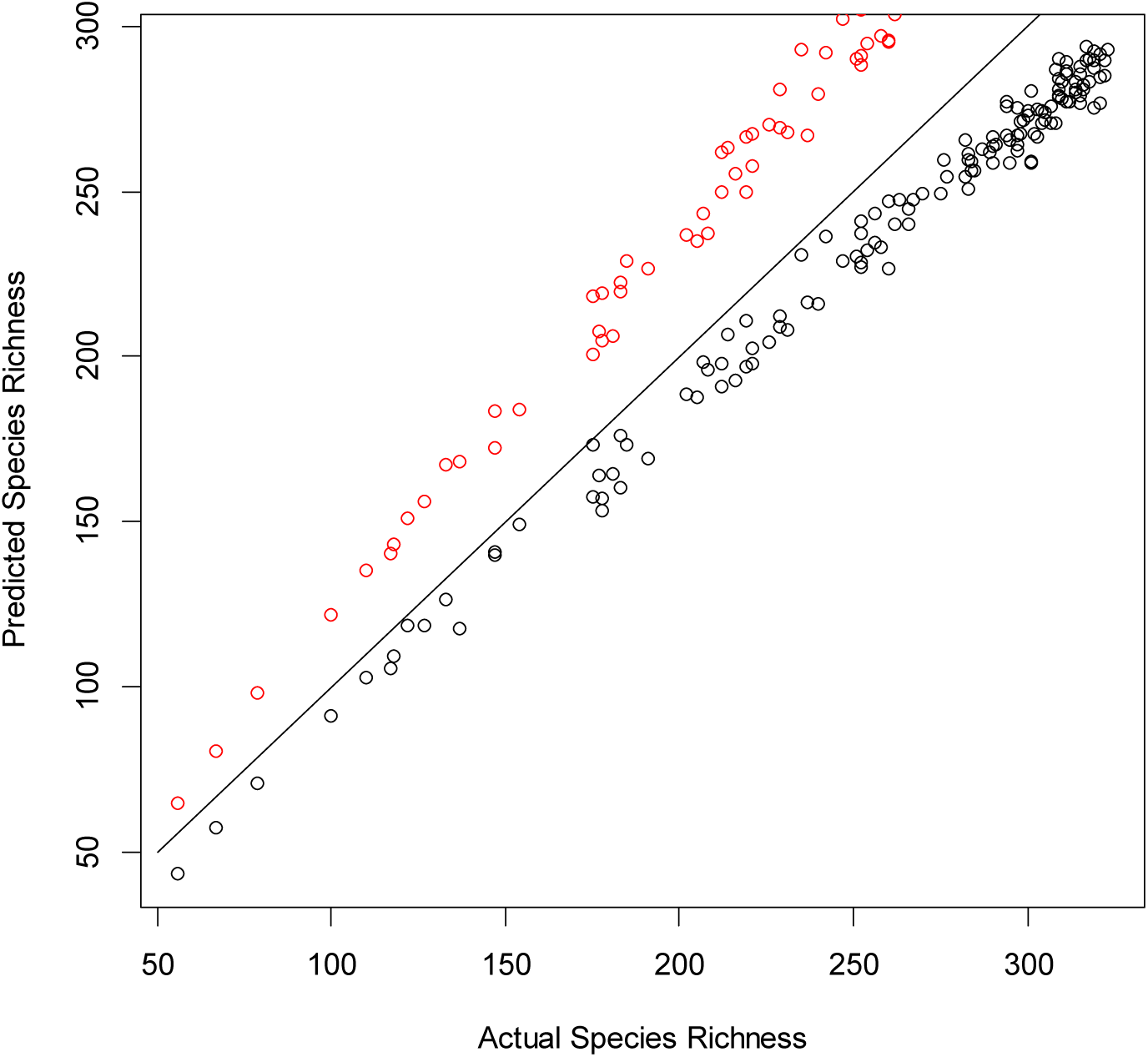
Estimated species richness at the end of each simulation, in function of the actual remaining species richness of that simulation after fifty time steps. Points above the diagonal where true value and estimate are equal are from calculations that include estimated local colonization effects. The points below the diagonal estimate species richness on the basis of local survival only.

### Checking for the occurrence of temporary emigration

The main text states that the models for detection probability suggest a limited presence of temporary emigration by entire species. Here I present further investigation of the issue.

It is implausible to assume that there would have been no temporary emigration of individuals in this study at all. Temporary emigration will affect estimation of species richness trends only if species have disappeared entirely from the study area for a substantial period, and have returned afterwards, and in such a way that the probability of temporary emigration was larger for a fraction of the time period. I simulated such temporary emigration of 10% of the species present between years 20 and 30 of a simulation running over fifty years (simulation code provided as a text file). A model with time trends in local survival and constant colonization was fitted to the resulting data, and detection probability either with categorical effects of time or a regression of covariates as in the main text. The differences between detection probabilities predicted by the two models were calculated (Figure S5, non-Bombus) and limits of confidence intervals compared. For the *Bombus* genus, we could basically not detect the temporary emigration. For non-*Bombus*, the time pattern of differences between the time-dependent and regression models indicates some detectability of the temporary emigration, but differences are also negative outside of the time window where it occurred. Between ten and twenty years in the simulation, 20% of the confidence intervals of the predictions are non-overlapping and in the direction we expect in the presence of temporary emigration (difference lower limit confidence interval prediction regression model and upper limit from prediction time-dependent model above zero). Between years twenty and thirty up to 46% of the predicted detection probabilities are significantly different (Fig. S5).

**Figure S5.**
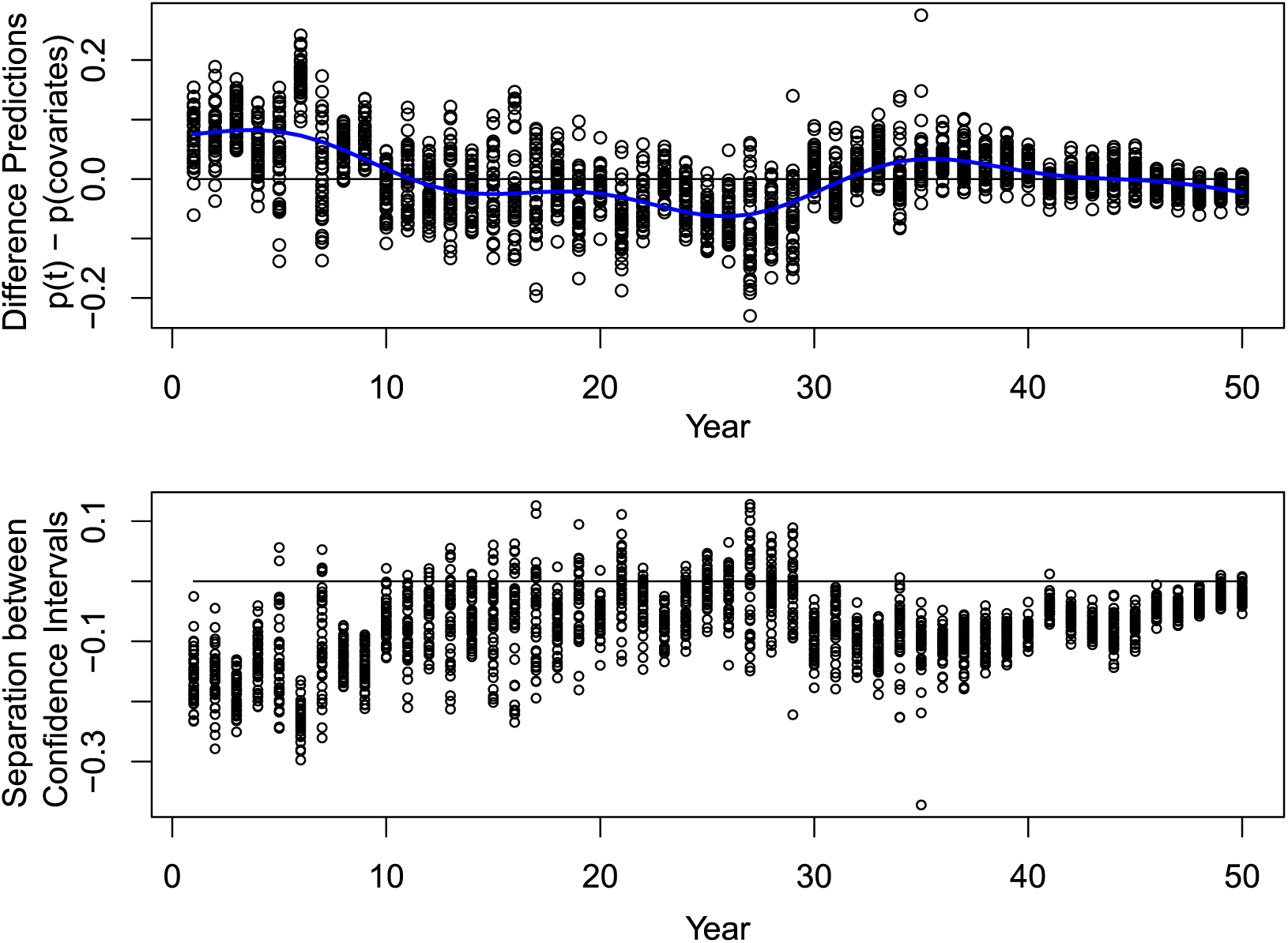
Results of simulations based on non-*Bombus* relative abundances where temporary emigration occurred for 10% of the species between years 20 and 30. First panel: differences per year between predicted detection probabilities of the categorical time-dependent model and the regression model. A generalized additive model with an automatic smoother estimation and selection procedure (method “GCV-Cp”; Wood 2006) was fitted to the differences and is drawn as a blue line. Bottom panel: separation between confidence intervals of predicted detection probabilities in the regression model and the categorical time-dependent model. When this separation is positive, the detection probability is significantly larger for the regression model, hence the categorical time effect is reduced which can indicate temporary emigration. Please note that positive separation values are concentrated between years ten and thirty.

